# SnapFISH-IMPUTE: an imputation method for multiplexed DNA FISH data

**DOI:** 10.1101/2024.01.12.575427

**Authors:** Hongyu Yu, Daiqing Wu, Guning Shen, Ming Hu, Yun Li

## Abstract

Chromatin spatial organization plays a crucial role in gene regulation. Recently developed and prospering multiplexed DNA FISH technologies enable direct visualization of chromatin conformation in nucleus. However, incomplete data caused by limited detection efficiency can substantially complicate and impair downstream analysis. Here, we present SnapFISH-IMPUTE that imputes missing values in multiplexed DNA FISH data. Analysis on multiple published datasets shows that the proposed method preserves the distribution of pairwise distances between imaging loci, and the imputed chromatin conformations are indistinguishable from the observed conformations. Additionally, imputation greatly improves downstream analyses such as identifying enhancer-promoter loops and clustering cells into distinct cell types. SnapFISH-IMPUTE is freely available at https://github.com/hyuyu104/SnapFISH-IMPUTE.

## Introduction

Chromatin conformation capture (3C) methods and their derivatives have been widely used for studying three-dimensional (3D) chromatin conformations.^1^ Sub-nuclear structures such as chromosome territories, topologically associating domains (TADs), and enhancer-promoter loops can all be identified from data generated by 3C-based methods.^2,3^ However, since 3C-based methods depend on crosslinking DNA fragments, downstream analyses often start from the pairwise contact matrices, and reconstructing 3D models from these matrices is challenging.^4–6^ Recent developments in fluorescence in situ hybridization (FISH)-based technologies, such as DNA multiplexed error-robust FISH (MERFISH)^7^ and DNA seqFISH+^8^, in contrast, do not rely on proximity ligation. Instead, a set of DNA probes is designed to cross-hybridize the genomic region of interest. By sequentially binding DNA probes with different fluorescence signals, the identity of each probe can be resolved, thus enabling direct visualization of DNA fragments in each single cell.^9^ However, the quality of FISH-based data decreases as the number of hybridization rounds increases. As a result, the average detection efficiency is below 80% for most chromatin conformation datasets with a resolution higher than 30Kb.^7,8,10–13^ Compared to 3C-based data, which only have dropouts represented by zeros in their contact matrices, FISH-based data have unobserved 3D coordinates that cannot be simply filled by a single value such as zero. This missing data problem makes many existing algorithms inapplicable. For example, none of the classical dimension-reduction algorithms for visualizations such as principal component analysis (PCA) or uniform manifold approximation and projection (UMAP) can handle missing values. To account for such missingness, two types of imputation strategies are adopted in published literature.^8,14^ The first approach imputes the pairwise distances between unobserved imaging loci. A missing entry in the distance matrix is imputed either by the average of neighboring entries^8^ or the average of that entry across all cells (mean imputation).^15^ This approach is appropriate if downstream analyses only depend on pairwise distances; otherwise, the problem of missing data pertains since the 3D coordinates of the unobserved loci are still missing. The second approach imputes 3D coordinates directly. The most common strategy is to perform linear interpolation by calculating a weighted average of two neighboring loci.^11^ Other methods such as nearest neighbors^16^ are also reported, but all existing strategies that impute 3D coordinates of missing loci rely on neighboring loci only and lack the ability to infer global structures. In practice, multiplexed DNA FISH data are typically filtered before analysis, leaving only the cells with detection efficiency higher than some predetermined threshold^8,16^, which further reduces the number of usable cells and complicates downstream analyses.

In contrast to the lack of imputation methods developed specifically for multiplexed DNA FISH data, computational methods tailored for imputing single-cell Hi-C (scHi-C) data, such as scHiCluster^17^ and Higashi^18^, have been proposed recently. However, these methods are not directly applicable to multiplexed DNA FISH data because they all build on assumptions made on 3C-data. For instance, the data are sparse at the single-cell level, measurements are discrete, and only dropouts are recorded instead of unavailable observations. To fill in such methodological gap, we present an imputation method tailored for multiplexed DNA FISH data in this paper. The imputation method relies on a dissimilarity measure to find chromosomes with similar conformations. Note by default, we perform analyses separately the two chromosomes in each cell. Then a target pairwise distance matrix is constructed for each chromosome based on the selected structures. To recover missing 3D coordinates, we minimize the difference between the target matrix and the spatial distances calculated from 3D coordinates. Compared to existing methods, our method is able to integrate information from both the neighboring loci of the missing locus and other cells. We have shown that the imputed data resemble real imaging data and can improve the results of various downstream analyses, including loop-calling and clustering.

## Results

### SnapFISH-IMPUTE

Multiplexed DNA FISH data measure the three-dimensional Euclidean coordinates of consecutive loci on chromosomes. Depending on the ploidy and the cell cycle, it is possible to observe multiple signals of the same locus within each cell. Here we assume that the identity of each locus, that is, to which allele it belongs, is already resolved by clustering or more advanced methods tailored for spatial genome alignment.^8,19^ Then the goal of imputation is to fill in the 3D coordinates of the missing loci on each chromosome.

Although chromosome conformations often demonstrate large cell-to-cell variability, common structures such as TADs and enhancer-promoter loops are conserved in subpopulations of cells from the same cell type.^8,10,11^ We therefore reason that for each missing locus, its position is not only related to loci that immediately upstream loic or downstream loci on the same chromosome but also can be determined by the relative positioning of the same locus on other cells. To find cells with similar structures, we first convert 3D coordinates to pairwise Euclidean distances between each locus pair. The pairwise distances are then normalized by their one-dimensional (1D) genomic distances, so the difference between two cells is not dominated by pairwise distance entries with large variations. Specifically, for locus pairs separated by the same 1D genomic distance, we model the difference in each coordinate as a centered normal distribution with variance proportional to the 1D distance. The squared pairwise distance, which is the sum of squares of the difference in each axis, thus follows a chi-squared distribution with three degrees of freedom multiplied by the variance. In imaging experiments, however, factors such as the resolution of the microscope and the imaging procedure used might lead to violations of the assumptions. To account for potential deviations, an additional Box-Cox transformation is performed, ensuring that the underlying distributions are approximately normal (Figure S6). Last but not least, we applied the z-score normalization to correct for the difference in 1D genomic distances (Figure 1).

**Figure 1.**
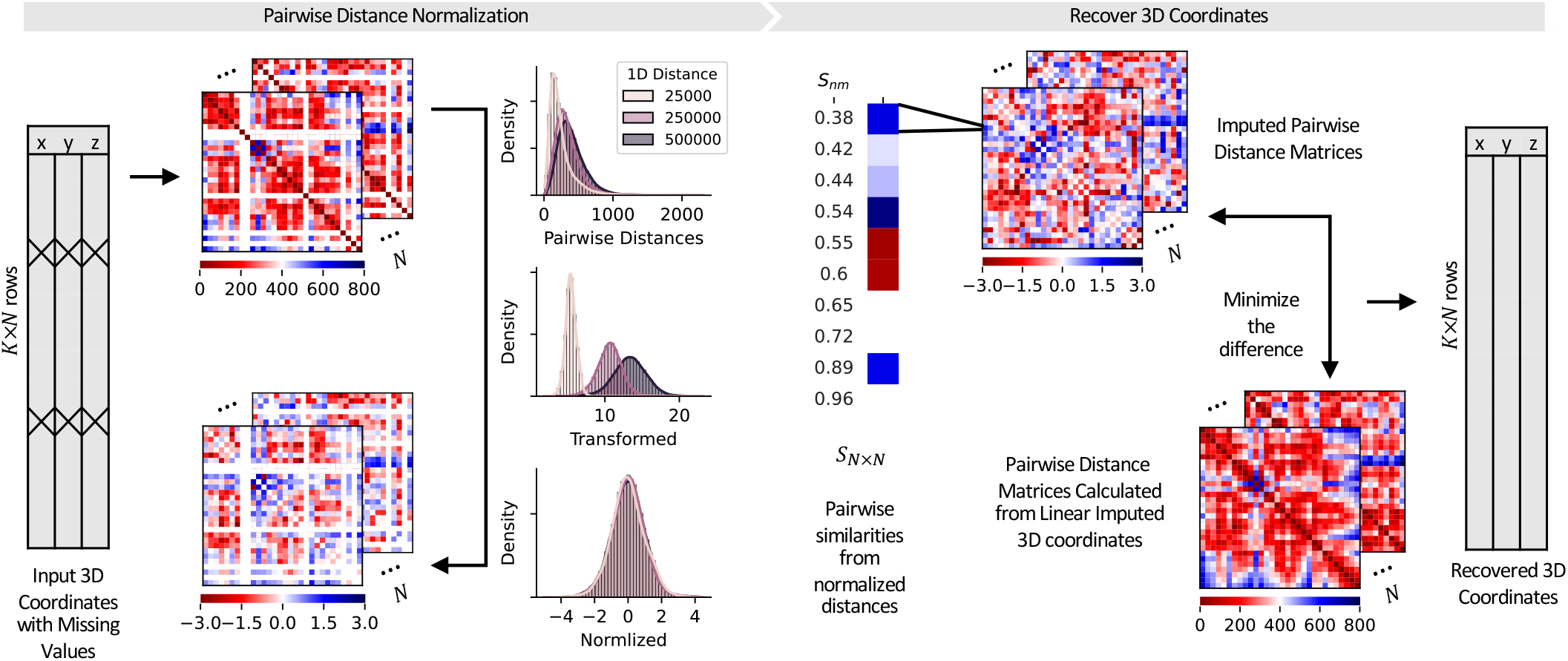
Overview of the imputation algorithm. The input includes the 3D coordinates of available loci in the imaging region and a 1D genomic location annotation file. The Euclidean coordinates are converted to pairwise distances and then grouped by 1D genomic distances. Our method adopts a two-step normalization procedure to systematically remove 1D genomic distance effects. The normalized distances are used to calculate dissimilarities between cells, which then determines how missing entries are filled. Finally, the 3D coordinates are recovered by minimizing the difference between the imputed pairwise distances and the distances calculated from linear initialized coordinates.

To find cells with similar conformations, we define a dissimilarity measure by computing the root mean square deviation between the normalized distances. Because of the presence of missing loci, when calculating the dissimilarity score between two cells, normalized distances that are unavailable in at least one of the two cells are skipped, and the final score is rescaled by the number of shared available entries, thus ensuring that all scores are within the same range and comparable with each other. We further filter the scores by removing the ones with shared entries less than 80% of the available entries in both cells. However, since the Euclidean distances are highly dynamic at the locus pair level, the dissimilarity defined in this way cannot capture larger structural features, such as TADs, a phenomenon also observed in Hi-C data.^20^ A widely-adopted solution for Hi-C data is to first smooth the contact matrix. Here, we resize the normalized distance matrices to 20 by 20 to achieve the same purpose. An additional advantage is that resizing will simultaneously reduce the missing ratio in the processed pairwise distances (Figure S1), hence allowing more scores to pass the 80% filter.

The next step of the imputation workflow involves constructing target pairwise distances. Specifically, for each cell, all the other cells are ranked by their dissimilarity, and each missing pairwise distance of the cell is replaced by the first non-missing entry in the sorted cell list. This replacing process is repeated for all cells until no missing pairwise distances exist. To recover the underlying 3D coordinates from pairwise distances, we minimize the difference between the target pairwise distances and the pairwise distances calculated from 3D coordinates. Similarly, the difference is calculated after pairwise distances are normalized by 1D genomic distances, which ensures that each entry contributes equally. The difference and its derivatives are then passed to a minimizer and optimized with the limited-memory Broyden–Fletcher–Goldfarb–Shanno algorithm (L-BFGS),^21^ a method designed for solving large-scale optimization problems.

### Imputation preserves the distribution of pairwise spatial distances

We applied our imputation algorithm to a published multiplexed DNA FISH dataset from mouse embryonic stem cells (mESCs).^10^ The authors performed a whole-genome DNA seqFISH+ on 446 cells from two biological replicates. Two groups of probes were designed, where the first group consists of 1,200 probes separated by 25kb and the second group consists of 2,460 probes separated by about 1Mb. We started with the 25kb resolution subset, where one imaging region is selected from each chromosome, and 60 consecutive loci separated by 25kb are imaged for each imaging region. Keeping only haploid chromatins with at least one locus observed, the average detection efficiency of the 25kb subset is 67.9% (range: 58.7% [chr2] 73.0% [chr3]). To benchmark our method SnapFISH-IMPUTE, we included three alternative imputation strategies in the following analyses: linear imputation, cubic spline imputation, and mean imputation (see Methods). The first two methods can recover the 3D coordinates while the mean imputation method can only fill in missing pairwise distances.

As expected, the average pairwise distances from the raw data and the mean imputed data are identical since in mean imputation, missing values are replaced by the average (Figure 2a). Linear imputation also gives an average distance matrix similar to the original one, though the distances between locus pairs with larger 1D genomic distance are more likely to be overestimated (Figure 2a, S2a). Interestingly, cubic spline, a generalization of linear imputation that uses a cubic polynomial instead of a linear function, completely removes the original patterns in the data (Figure 2a, S2b). This suggests that filling in missing 3D coordinates while preserving population-level features is challenging. The main reason that linear imputation works reasonably well is likely that arithmetically, it is equivalent to averaging neighboring two loci, so it implicitly captures the stochasticity in chromatin conformation and replaces those missing coordinates by their expectations. If we try to capture the variations explicitly by increasing the order of the fitted function, the predicted value is no longer computed from the neighboring two loci only but will also be influenced by other loci further apart. In contrast, by adopting a two-stage imputation workflow, SnapFISH-IMPUTE preserves the original population-level patterns faithfully and yields almost identical distance matrices (Figure 2a, S2c). The difference between the imputed matrix and the raw matrix is also substantially smaller than the difference computed from both the linear imputation result and the cubic spline imputation result (Figure 2b).

**Figure 2.**
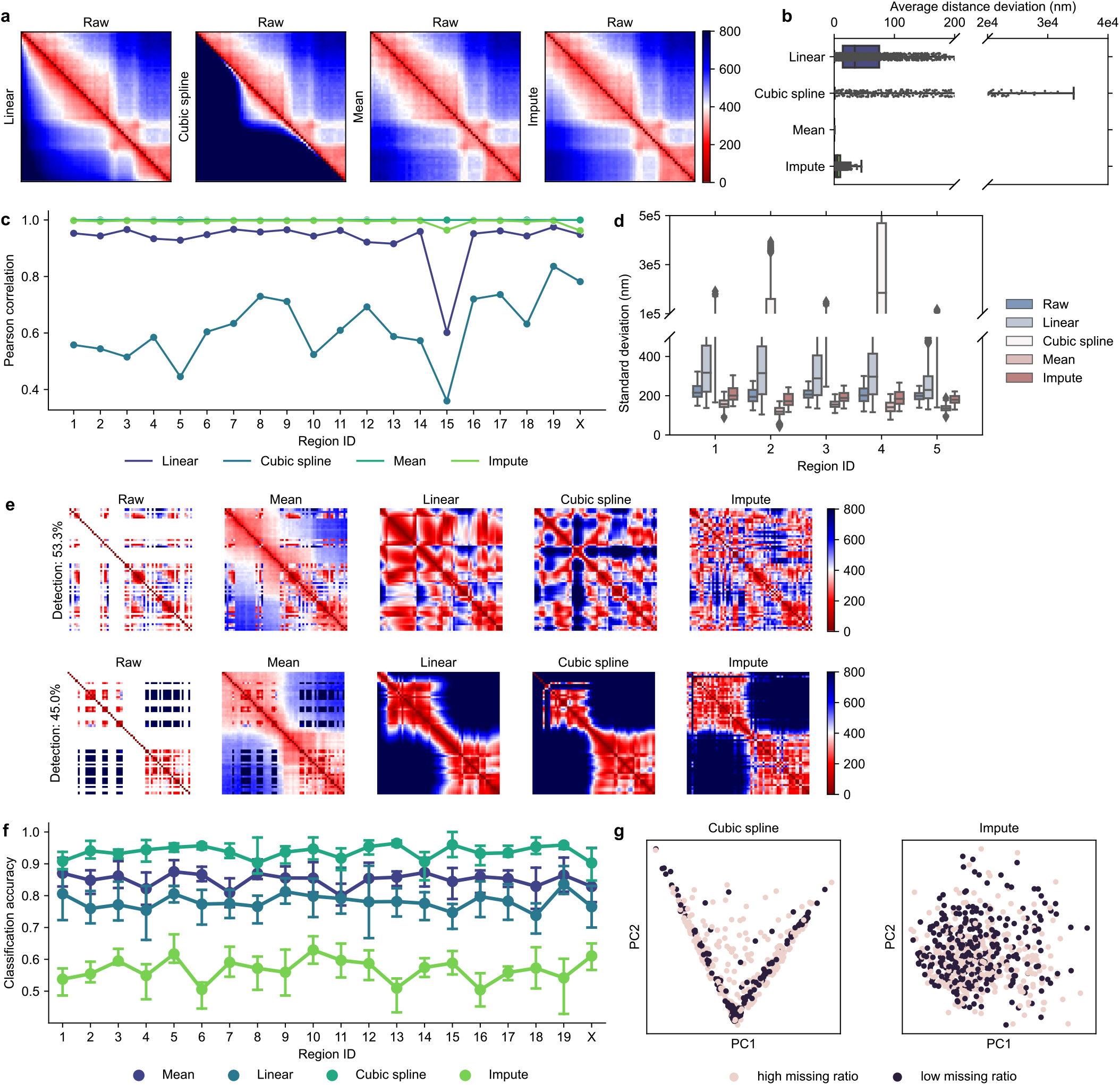
Imputation results of the 25kb subset of the DNA seqFISH+ mESCs dataset. **a** The average pairwise distance matrices of imaging region 1. The upper triangle is the raw average distance matrix, and the lower triangle is the imputed distance matrix. **b** The absolute value of the difference between the upper and the lower triangle in part **a. c** The Pearson correlations between the average distance matrices from the raw data and the imputed data across the 20 imaging regions. **d** The standard deviations of each entry in the pairwise distance matrix. The first five imaging regions are shown (see fig S2 for the other 15 regions). **e** Single cell examples. **f** Binary classification accuracies across different imaging regions. **g** The first two principal components of mean imputation result and our imputation method. The groups with the highest and the lowest missing ratios from imaging region 4 are shown.

The Pearson correlations between the mean distance matrices from the imputed data and the raw data show that SnapFISH-IMPUTE consistently outperforms linear imputation and cubic spline across all imaging regions. Indeed, the correlations are around 0.99, almost the same as the ones from mean imputation (Figure 2c). Such high correlations are achieved without gathering population-level information beforehand, which is completely different from mean imputation, where the population average is pre-computed and directly used to fill in missing pairwise distances. The results indicate that the information from other cells is referenced sufficiently in the first part of our method, and the 3D coordinates recovered resemble true distributions. In addition to the mean distance matrix, we calculated the standard deviation of the distance between each locus pair across the dataset. Both linear imputation and spline imputation lead to unwanted variations, which can be more than twice as high as the observed variations (Figure 2d). The mean imputation, on the other hand, reduces the original standard deviations because all missing distances from the same locus pair are replaced by the same value. Among all methods tested, although our proposed imputation method also slightly decreases the variation, it keeps the overall distribution at a similar level (Figure 2d, S2d). Taken together, our method preserves population-level features and surpasses other benchmark methods in multiple aspects.

### Imputed conformations resemble observed conformations

We next evaluated the effect of imputation on single-cell level characteristics. A few cells without missing loci are selected, and we found that chromatin conformations demonstrate large cell-to-cell variability, coherent with previous studies (Figure S3a).^10^ Nevertheless, there are some common patterns shared by most imaged regions. For example, the loci are often arranged sinuously instead of following a smooth curve. As a result, neighboring entries in the pairwise distance matrix can have distinctive values, despite their being close in 1D genomic distance (Figure S3a). Such features are not always preserved by other imputation methods considered. When about 50% loci are missing, both linear imputation and cubic spline imputation fail to recover the sharper changes observed in real imaging data (Figure 2e). This pattern is more obvious as the detection efficiency drops below 50%, where loci start to pack together and finer structural variations are entirely lost, as reflected by the distance matrices (Figure S3e,f). SnapFISH-IMPUTE, on the opposite, is robust under various detection efficiencies, and the imputed loci distribute randomly as in real imaging data even when two-thirds of the loci are missing (Figure S3b-f). Remarkably, although SnapFISH-IMPTUE does not model the conformation parametrically, it captures the overall trend of non-missing loci, and the imputed pairwise distances are indistinguishable from the real pairwise distances, in contrast to mean imputation (Figure 2e).

If the imputed loci emulate real imaging data at the single-cell level, it would be difficult to discriminate imputed cells with low detection efficiency from those with high detection efficiency. For each imaging region in the 25kb subset, we binned the cells by their detection efficiencies into three groups, and we trained a soft-margin support vector machine on the two groups with the highest and the lowest detection efficiencies (see Methods). If the imputed result closely resembles true data patterns, the classifier would not be able to distinguish cells from these two groups easily. Indeed, the accuracy is around 50% for SnapFISH-IMPTUE across different imaging regions, which is about the same as a random classifier. In comparison, the classification accuracies are considerably higher for all three competing methods, with mean imputation having the highest score of around 90% (Figure 2f). We performed a principal component analysis (PCA) using the mean imputed data and the data generated by SnapFISH-IMPUTE. The result shows that cells with low and high detection efficiencies from the mean imputed data occupy different sub-spaces in the low-dimensional space, consistent with the high classification accuracies. In contrast, cells with different detection efficiencies are mixed together in the PCA plot of SnapFISH-IMPUTE, confirming that the predicted conformations are similar to the observed ones (Figure 2g).

### SnapFISH-IMPUTE is robust under different resolutions and imaging protocols

Next, we applied SnapFISH-IMPUTE to the 1Mb subset of the mESCs data. We imputed missing data in output from Jie^19^, a spatial genome alignment method. The overall detection efficiency is only 36.3%, with range 30.2% 46.9%. Notably, although nearly two-thirds of the 3D coordinates are missing, SnapFISH-IMPUTE yields average distance matrices almost identical to the original ones, and the Pearson correlations are close to one as before (Figure S4a,b). Additionally, we have analyzed a DNA FISH dataset of mESCs generated by a different imaging protocol.^11^ A total of 41 probes are designed to image two alleles at 5kb resolution, and the average detection efficiency is about 70% for both alleles. Similar to previous results, SnapFISH-IMPUTE recovers missing 3D coordinates effectively, as reflected by both the mean distance matrices and the distance deviations (Figure S4c-f). The correlations are 0.98 and 0.98 for the two alleles, in contrast to the 0.88 and 0.86 achieved by linear imputation and the 0.58 and 0.57 by cubic spline imputation. In summary, the performance of the SnapFISH-IMPUTE is robust under different imaging resolutions and protocols.

### Imputation improves downstream analysis

Enhancer-promoter loop is a critical structural feature in chromatin conformation data and plays a key role in transcription regulation.^22^ In our recent work, we have developed a loop-caller, SnapFISH^23^, for multiplexed DNA FISH data. We applied SnapFISH to the 25kb imputed DNA seqFISH+ data and benchmarked the loop-calling performance with both the loop set from the raw data and the HiCCUPS^24^ output. With the default threshold, SnapFISH identified 30 loops from the imputed data, of which 14 loops overlapped with the HiCCUPS output (Figure 3a). The large number of false positives is not surprising since imputation increases the effective sample size and thus allows more loops to pass the default threshold. The *t*-test results from SnapFISH show that false positive loops have smaller *t*-statistics than true positive loops, suggesting that the relative loop strength is not affected by imputation (Figure 3c). This motivates us to optimize the default FDR cutoff in the SnapFISH algorithm to achieve a similar precision to the raw data. Indeed, we found that by setting the cutoff to 0.001, SnapFISH reports 4 false loops while all the true loops are not affected (Figure 3b). It is worth noting that not all imputation strategies will enhance loop-calling. Both linear imputation and cubic spline imputation significantly lower the sensitivity, with only two loops called at the optimal threshold. We reason that enhancer-promoter loop is a more intricate feature in 3D chromatin conformation, thus requiring careful processing of the raw data. Next, we applied SnapFISH to the 5kb chromatin tracing data and tested whether it could identify the *Sox2* enhancer-promoter loop with different numbers of cells. Specifically, we generated 100 random samples for each number of cells and calculated the F1 score on these 100 samples. The result shows that imputation boosts loop-calling efficiencies and leads to a higher F1 score across both alleles (Figure 3d,e).

**Figure 3.**
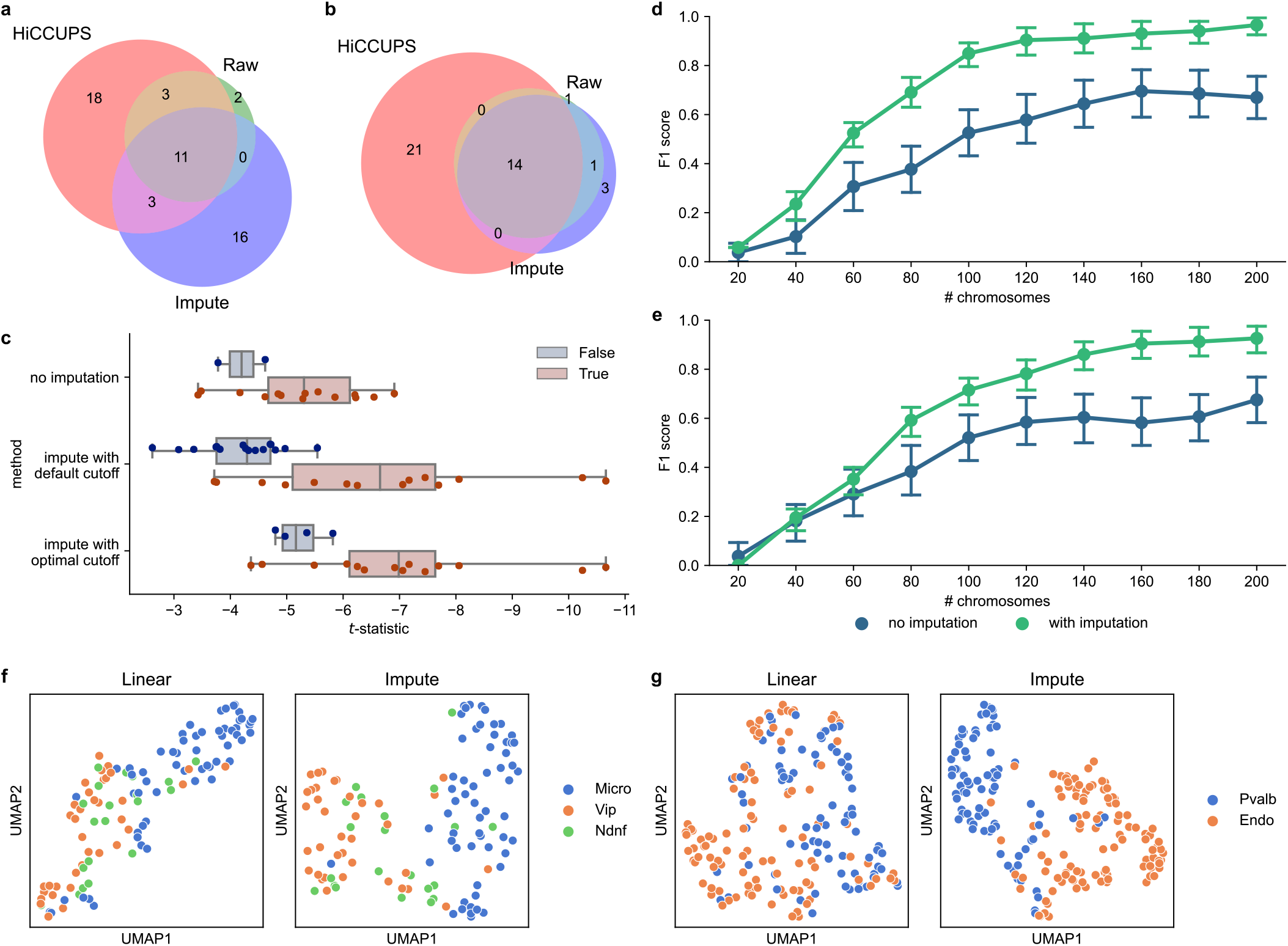
Imputation improves loop-calling and cell type clustering. **a** Loops called from the mESCs 25kb DNA seqFISH+ dataset using the default threshold. **b** Loops called from the 25kb subset using the optimal threshold in SnapFISH. **c** The *t*-statistics of loops called by SnapFISH. **d** Identification of the Sox2 enhancer-promoter interaction on the 129 allele. **e** Identification of the Sox2 enhancer-promoter interaction on the CAST allele. **f g** The embedding of different mouse brain cell types.

In addition to loop-calling, we test whether the imputed data can be used for cell-type clustering. We re-analyzed a previously published DNA seqFISH+ dataset of mouse brain cells^8^. The authors also conducted mRNA seqFISH+ on each cell in the dataset and identified nine major cell types. Here we asked whether cells have distance 3D conformations in different cell types. We applied SnapFISH-IMPUTE and linear imputation to the 1Mb resolution subset of the data. After obtaining the complete 3D coordinates, we computed the pairwise distances of each cell and concatenated all pairwise distances from the 19 autosomes in each cell. Since not all cells have both alleles observed and all 19 autosomes recorded, we kept only cells with at least one haploid observable and randomly select one allele for cells with more than one allele to perform clustering. The concatenated distances are normalized and then embedded with PCA followed by UMAP. Although not all cell types are distinguishable from each other in the embedding space, some of them, such as microglia (Micro) and neurons expressing vasoactive intestinal polypeptide (Vip) (Figure 3f) or neurons expressing parvalbumin (Pvalb) and endothelial cells (Figure 3g), occupy distinct regions. We also noticed that SnapFISH-IMPUTE often separates these classes more clearly than linear imputation and thus is more appropriate for downstream analyses.

To quantitatively evaluate the clustering efficiency, we calculated the adjusted mutual information scores (AMI) between the imputed data and the ground truth. Specifically, we first embedded the distances with PCA and then applied hierarchical clustering to obtain the predicted cluster assignments. We then computed the AMI score using the predicted and the true cluster assignments. The AMI scores of SnapFISH-IMPUTE are almost always higher than the ones from linear imputation (Figure S5). For example, for pairs Pvalb versus Endo, Micro versus Vip, and Sst versus Micro, linear imputation yields an AMI close to zero, indicating that the clusters found are almost random. In contrast, SnapFISH-IMPUTE give an AMI scores close to 0.5 in these cases, which are considerably higher and much closer to 1, showing that SnapFISH-IMPUTE preserves original chromatin conformation characteristics.

## Discussion

In this paper, we presented SnapFISH-IMPUTE, a unified nonparametric method to impute missing values in multiplexed DNA FISH data. In contrast to previous attempts, which focused on either imputing 3D coordinates with simple parametric models or developing ad-hoc methods for specific purposes, our method takes into account that cells imaged together have common structures. By allowing similar cells to borrow information from each other, we improved the quality of the imputed data significantly. We systematically evaluated the performance of SnapFISH-IMPUTE on three published multiplexed DNA FISH data of mice under different resolutions and experiment protocols. The results showed that the spatial relationships between DNA loci are preserved in the imputed data, and a similar level of cell-to-cell variation is observed after imputation. By dividing the cells by their missing ratios and training a classifier on the imputed data, we have also demonstrated that SnapFISH-IMPUTE performs consistently regardless of missing ratios. In addition, we have benchmarked our method with three imputers used in previous studies and confirmed that each of them has limitations, introducing confounding variables that can influence following analyses.

More importantly, the imputed data outputted by SnapFISH-IMPUTE can be used for downstream analyses with minimal extra processing. For example, we showed that the imputed mouse brain cell data can be directly used for cell type clustering, with no additional adjustments needed. Furthermore, biological features became more detectable in the imputed data. As demonstrated with the 5kb chromatin tracing data, by increasing the effective sample size, imputation improves the F1 score of loop calling and thus makes it possible to identify enhancer-promoter interactions with fewer cells. We also found that the normalization procedure proposed in this work effectively removes biases and transforms the pairwise distances between DNA loci with a fixed 1D genomic distance to approximately normal (Figure S6f). In comparison, normalization by *z*-score only or by taking the Pearson residuals from generalized linear models cannot standardize the distributions effectively (Figure S6d,e).

We acknowledge that the efficiency of SnapFISH-IMPUTE might be compromised if the dataset contains a limited number of cells or if some subpopulations have distinct conformations but a small population size. In these cases, SnapFISH-IMPUTE might not be able to find cells with similar conformations and return imputed data that fail to represent the true biological relations.

In summary, SnapFISH-IMPUTE is an effective algorithm for imputing missing 3D coordinates in multiplexed DNA FISH data. The improvement in data quality through imputation will enable more computational and statistical tools to be developed, bringing new insights to the understanding of gene regulations in cells.

## Methods

### Pairwise Distance Normalization

Let (*x*_*in*_, *y*_*in*_, *z*_*in*_) denote the *x, y*, and *z*-coordinates of the *i*-th locus in the *n*-th cell. The Euclidean distance between the *i*-th and the *j*-th loci is defined by

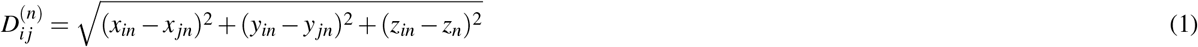

If we assume that the measurement error in each axis is the same, then their differences should be similar. Additionally, the distribution of *x*_*i*_ *-x* _*j*_, *y*_*i*_ *-y* _*j*_, and *z*_*i*_ *-z* _*j*_ should have a mean of 0; otherwise, it means that cells from different cells are all aligned in the same direction, which is unlikely. For locus pairs with the same 1D genomic distances, *x*_*i*_ *-x*_*j*_, *y*_*i*_ *-y*_*j*_, and *z*_*i*_ *-z*_*j*_ can be viewed as three independent centered normal distributions with the same variance. Therefore, after grouped by 1D distances, *D*^2^ divided by the common variance is the sum of squares of three standard normal distributions, thus following a 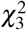 distribution. However, depending on the experiment design, the imaging data might be acquired in a slice-by-slice manner, leading to an unequal variance in each axis^10^ Additionally, the resolution limit of microscopes might lead to different measurement errors. As a result, although the distances distribute closely as chi-squared distributions, they may possess distinctive skewness and tailedness.

Since *D* is non-negative and roughly follows a chi-squared distribution, we can apply Box-Cox transformation to standardize it^25^. We determine the optimal parameter *λ* by maximizing the likelihood of the transformed distance:

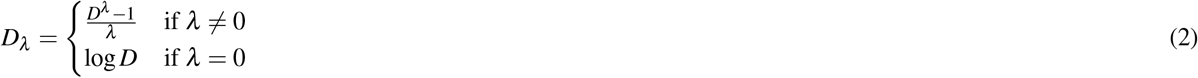

Then *D*_*λ*_ is normalized by (*D*_*λ*_ *-μ*)*/σ*, where *μ* and *σ* are the mean and the standard deviation of the transformed distances with the same 1D genomic distance. After normalization, we expect pairwise distances to distribute identically as standard normal distributions and contribute equally in the following analyses.

### Dissimilarities between cells

As noted in the HiCRep paper^20^, domain structures in contact maps are more stable features than individual interactions. Therefore, comparing larger structures instead of highly dynamic single interactions should be the main goal of a dissimilarity measure. Although HiCRep was developed for bulk Hi-C data comparison, the distance matrix calculated from multiplexed DNA FISH data is negatively correlated with Hi-C contact matrix, so similar issue might arise. Moreover, we are comparing single-cell level structures instead of aggregated means in this case, so each pair of conformations is more variable. To remove the noise, we resize the normalized distance matrix by a factor of *k* such that the resulting matrix has a side length just above 20. Let 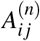 be the normalized pairwise distance between the *i*-th and the *j*-th loci of the *n*-th cell. Then the *i j*-th entry of the resized matrix *H*^(*n*)^ is the mean of the available entries in *A*^(*n*)^over the window of (*k*(*i -* 1), *ki*] *×* (*k*(*j -* 1), *k j*]. If the side length of *A*^(*n*)^ is not divisible by *k*, a zero padding is applied, which corresponds to the mean of a standard normal distribution.

The dissimilarity between the *n*-th and the *m*-th cell is then defined by

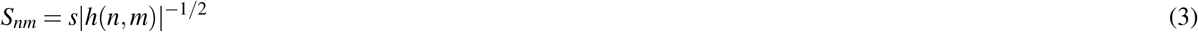

where |*h*(*n, m*)| is the number of entries where both *H*^(*n*)^ and *H*^(*m*)^ have values, and *s* is defined by

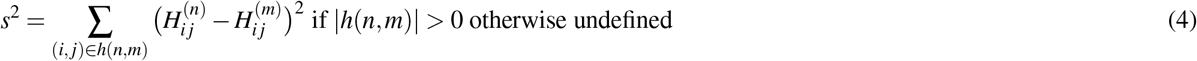

This is the Euclidean distance between the available entries weighted by the number of shared entries. If only a small proportion of the observed pairwise distances are shared between the two cells, the value defined might not reflect the true structural relation. Therefore, we further filtered the dissimilarities by the number of shared entries. Specifically, we calculate

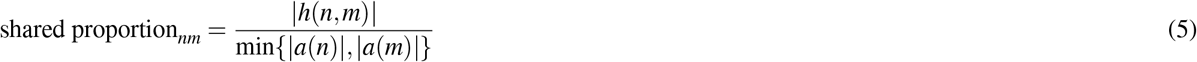

for each cell pair, where *a*(*n*) and *a*(*m*) are the sets of non-missing pairwise distances in cell *n* and cell *m. S*_*nm*_’s with ratio below 0.8 are left undefined.

### Impute missing pairwise distances

For the *n*-th cell, let *p*(*n*) be the set of indices where *S*_*nm*_, *m ∈ p*(*n*) is defined. For the missing locus pair (*i, j*) of cell *n*, define *u*(*i, j*) be the indices of all cells where 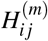, *m ∈ u*(*i, j*) is available. Thus, the set *p*(*n*) *∩u*(*i, j*) is the indices of all cells with the (*i, j*)-th entry available and has a dissimilarity score with cell *n*. We replace the missing pairwise distance (*i, j*) of cell *n* with

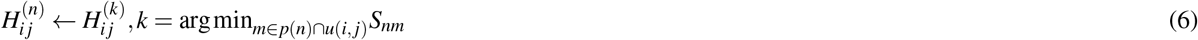

It is possible that *p*(*n*) *∩u*(*i, j*) is empty as no *S*_*nm*_ is defined or no *m ∈ p*(*n*) has entry (*i, j*). In such cases, we impute as many pairwise distances as possible and use the updated distances to recalculate the dissimilarities. Then another round of imputation is performed to fill in more missing pairwise distances. Since the missingness decreases after one iteration, both *p*(*n*) and *u*(*i, j*) contain more indices. By repeating the replacement procedure, all previously unavailable pairwise distances will eventually be imputed.

### Recover 3D coordinates from pairwise distances

To recover the underlying 3D coordinates from pairwise distances, we first initialize the missing (*x*_*in*_, *y*_*in*_, *z*_*in*_) by linear imputation (see linear imputation and cubic spline imputation). Let 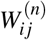 be the pairwise distance computed from the lineraly imputed data.

The same Box-Cox transformation and *z*-score normalization are used to transform 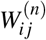 to 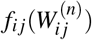 where

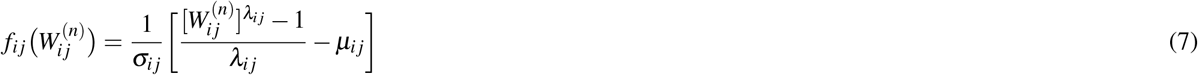

The difference between the target pairwise distance *H*^(*n*)^ and *f* (*W* ^(*n*)^) is then defined by

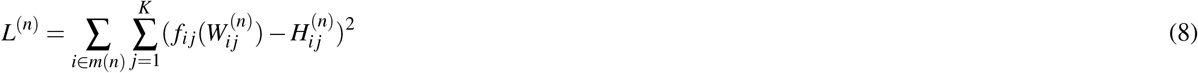

where *m*(*n*) is the set of missing locus on cell *n* and *K* is the total number of loci in the imaging region. The partial derivatives with respect to *x*_*in*_, *y*_*in*_, and *z*_*in*_ are

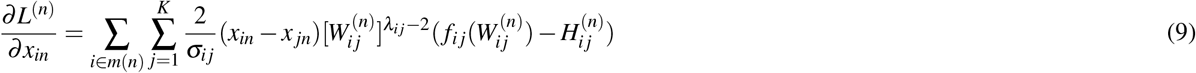

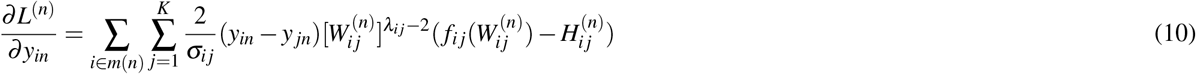

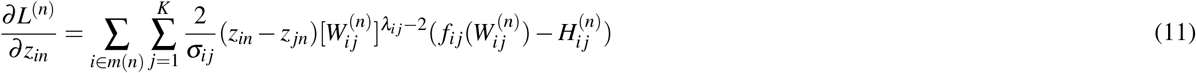

Both *L*^(*n*)^ and its partial derivatives are passed to the minimize function in SciPy^26^ and optimized with the L-BFGS algorithm, a quasi-Newton method that stores only a limited number of iterations to accelerate its convergence^21^. The optimized 3D coordinates are the imputation results.

### Time cost of SnapFISH-IMPUTE

SnapFISH-IMPUTE is implemented with mpi4py^28^ to allow multiprocessing. With 40 processes, SnapFISH-IMPUTE takes about 5 minutes to impute the 1,298 haploid chromosomes in the 5kb chromatin tracing data from Huang et al^11^. The time cost grows roughly linearly with the number of haploid chromosomes in the dataset.

### Linear imputation and cubic spline imputation

For each chromosome, a parametrized continuous spline is fitted for each axis separately:

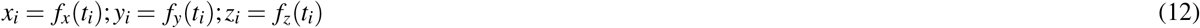

where *t*_*i*_ is the 1D genomic location of the *i*-th observed locus on the chromosome. For the *j*-th missing locus, its 3D coordinates are imputed by evaluating the fitted splines at genomic location *t* _*j*_ (i.e. 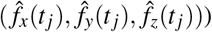. If *f* is a linear function, then this corresponds to linear imputation and can be interpreted as an weighted average of two neighboring loci.^11,14^ If *f* is a cubic polynomial, then this is the cubic spline imputation. Both methods are implemented with SciPy’s interp1d function from the interpolate module.^26^

### Mean imputation

As defined above, let 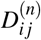 denote the Euclidean distance between locus *i* and locus *j* in chromosome *n*, and let *u*(*i, j*) denote the set of indices that 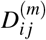, *m ∈ u*(*i, j*) is defined. We calculate the mean of each locus pair by

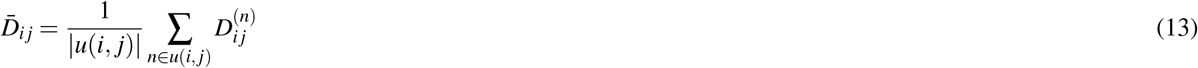

and replace each missing pairwise distance by

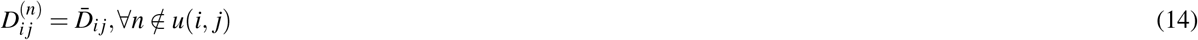

### Classify haploid chromosomes by detection efficiencies

For each imaging region, we first calculated the detection efficiency of each chromosome using the raw data. The chromosomes are then ranked and binned by the 33.3% and the 66.7% quantiles of detection efficiencies. We then calculated the pairwise distance matrices and normalize each entry of the distance matrix by its average across the dataset. We next divide the first and the third tertiles of the binned data by a five-fold cross validation. A soft-margin support vector machine with a radial kernel (implemented with the SVC class from the scikit-learn^27^ package) is then fitted to the training set, and the classification accuracy is calculated on the validation set. This training and validation procedure is carried out on each possible partition of the data.

### Statistics and reproducibility

No statistical tests were conducted to draw conclusions. No statistical methods were used to determine sample sizes. Unless otherwise noted, error bars correspond to 95% confidence intervals. No representative results were selected from repeated measurements. The proposed method was tested using three publically available datasets, and no specific observations were excluded.

## Data availability

The DNA seqFISH+ data of mouse embryonic stem cells (mESCs) are downloaded from Zenodo (https://zenodo.org/records/3735329). The 5kb chromatin tracing data are downloaded from 4DN Data Portal with ID 4DNESC5PKTQ9. The DNA seqFISH+ data of mouse brain cells are downloaded from Zenodo (https://zenodo.org/records/4708112). The aligned results from Jie are downloaded from 4DN Data Portal with IDs 4DNFIS6MLXGA and 4DNFIU73OR5W. The code for all data preprocessing is available at https://github.com/hyuyu104/SnapFISH-IMPUTE/blob/main/jupyter/preprocess.ipynb. The imputed data used in this study are available at https://zenodo.org/records/10088109.

## Code availability

The SnapFISH-IMPUTE software is freely available as a Python package at https://pypi.org/project/sfimpute. The documentaion is available at https://github.com/hyuyu104/SnapFISH-IMPUTE.

## Acknowledgements

This study was funded by the NIH grants R35HG011922 (to M.H.) and U01DA052713 (to Y.L.). Y.L. was also partially funded by the NIH grants U01HG011720, and U24AR076730. Ming Hu was also partially funded by the NIH grants UM1HG011585.

## Author contributions

H.Y. implemented the SnapFISH-IMPUTE software. Y.L. and M.H. supervised the project. H.Y., D. W., and G.S. performed data analysis and evaluated the method. H.Y., M.H., and Y.L. wrote the manuscript with input from all the authors. All authors read and approved the final manuscript.

## Competing interests

The authors declare no competing interests.

---

**Figure S1.**
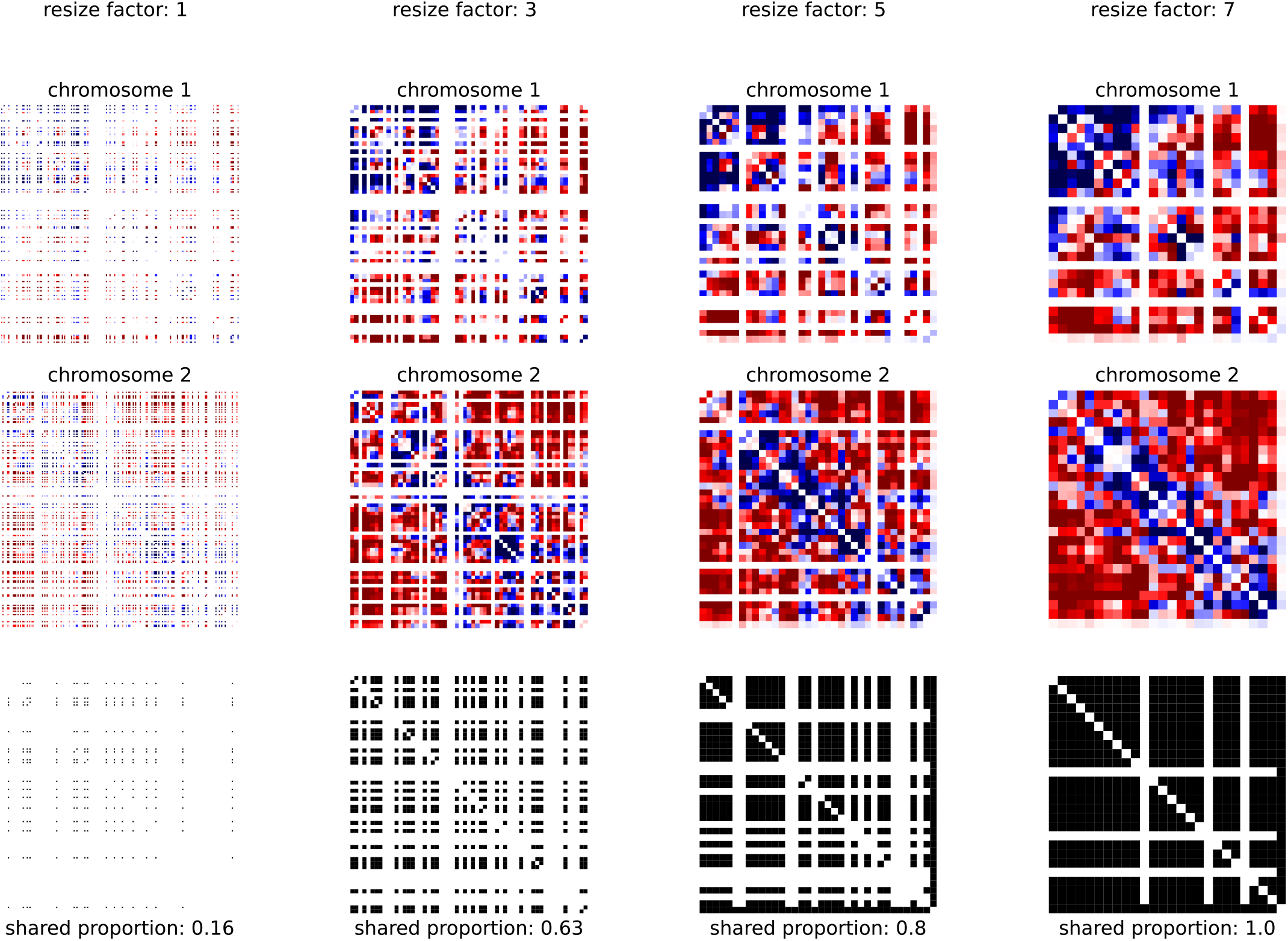
More overlapping values after resizing. The first row and the second row show the pairwise distance matrices of two cells under different resizing ratios. The last row shows the shared available entries between the two cells under different resizing ratios.

**Figure S2.**
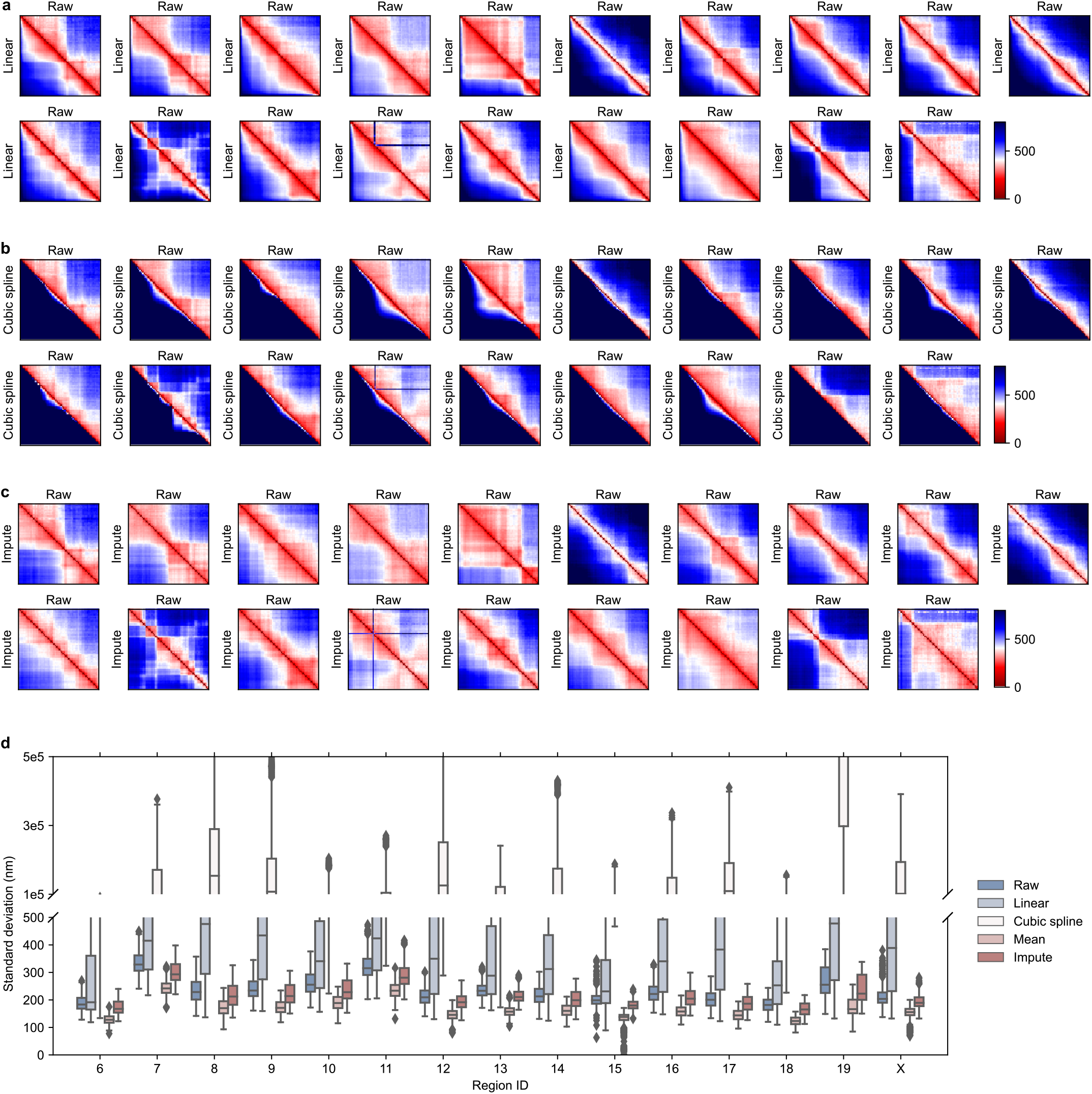
The averages and the standard deviations of distance matrices from the 25kb subset of the DNA seqFISH+ mESCs dataset. **a-c** Average pairwise distance matrices calculated from linear imputation, cubic spline imputation, and the proposed imputation method. Region 2 to 20 (autosome 2 to 19 and X cell) are shown. **d** The standard deviation of each entry in the distance matrix. Region 6 to 20 are shown.

**Figure S3.**
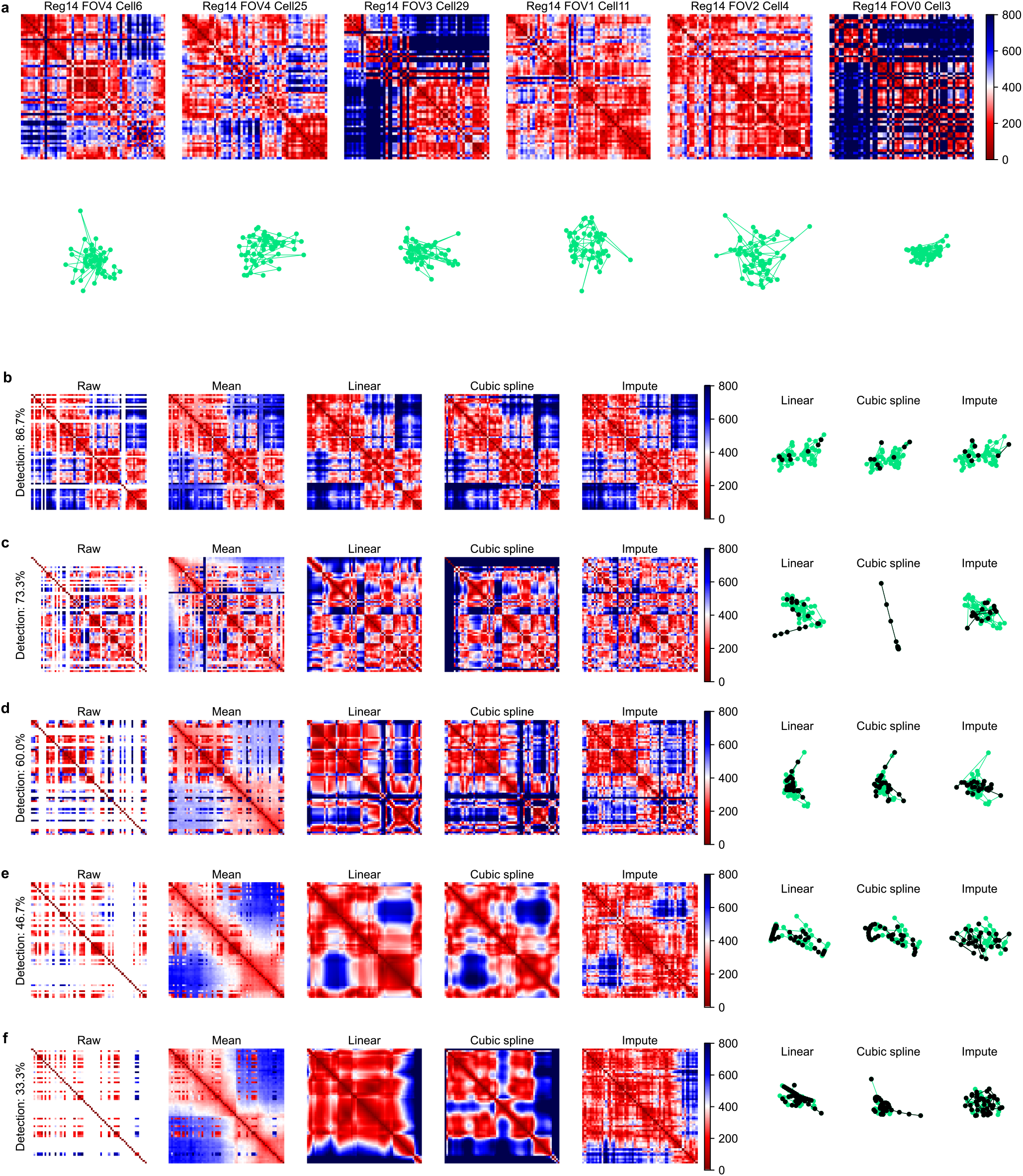
Single cell examples. **a** Cells with no missing loci are selected from the 25kb subset of the DNA seqFISH+ mESCs dataset (Reg: imaging region; FOV: field of view; Cell: cell ID). The first row is the pairwise distance matrices, and the second row is the 3D conformations. **b-f** Cells with decreasing detection efficiencies. The distance matrices of the raw data and the imputed data are shown. The last three plots in each line are the 3D conformations from the linear imputation result, cubic spline imputation result, and the imputation result from our method. Green dots are observed loci, and black dots are imputed loci.

**Figure S4.**
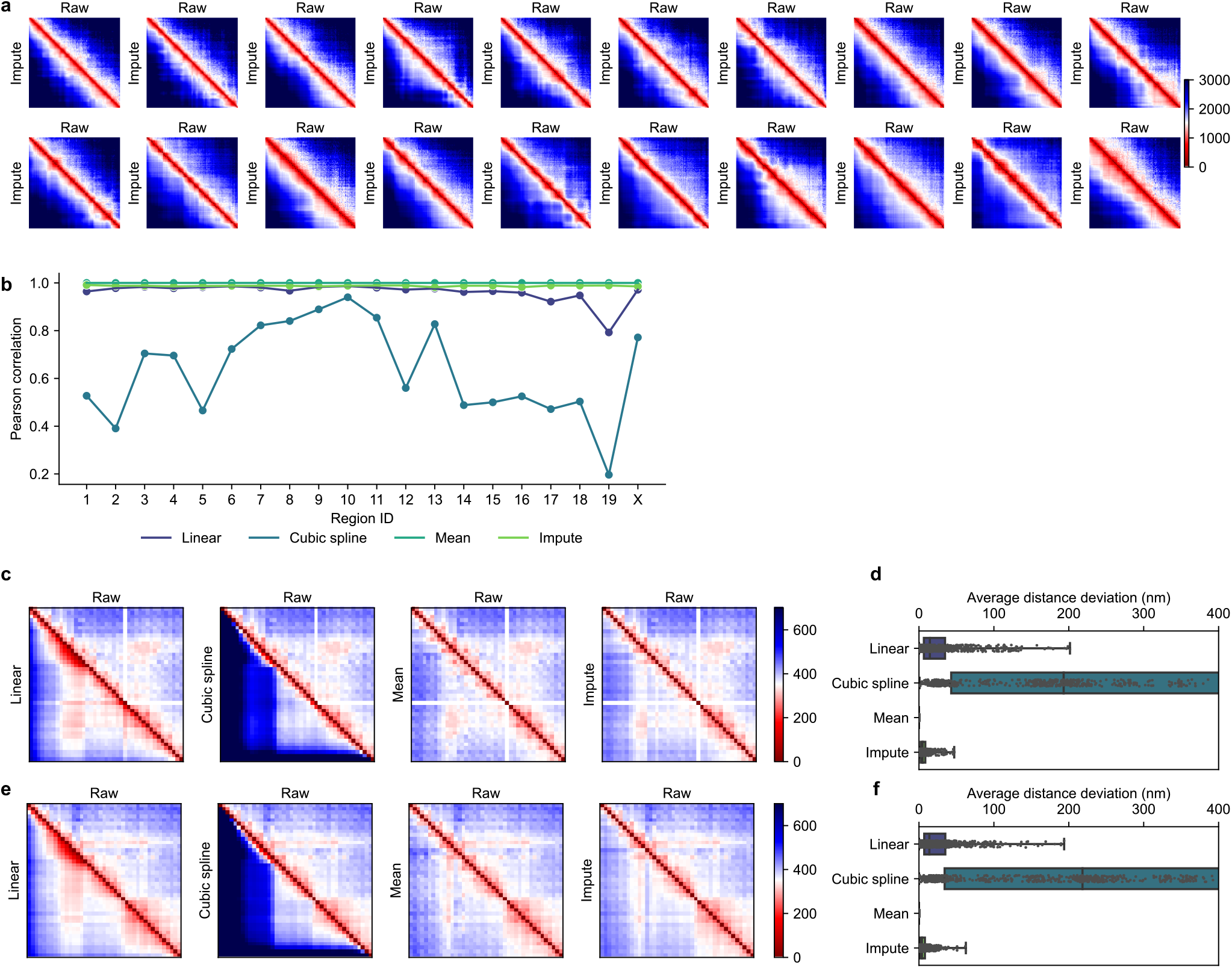
The proposed imputation method generalizes well to other imaging datasets with different resolutions and imaging protocols. **a** The average distance matrix of each imaging region in the 1Mb subset of the DNA seqFISH+ mESCs dataset. The upper triangle is the raw distance matrix, and the lower triangle is the imputed distance matrix. **b** Pearson correlations between the raw average distance matrix and the average distance matrix calculated from different imputation methods. **c** Average pairwise distance matrix of the 129 allele from the 5kb chromatin tracing dataset. **d** The absolute difference between the upper triangle and the lower triangle in part **c. e** Average pairwise distance matrix of the CAST allele from the 5kb chromatin tracing dataset. **f** The absolute difference between the upper triangle and the lower triangle in part **e**.

**Figure S5.**
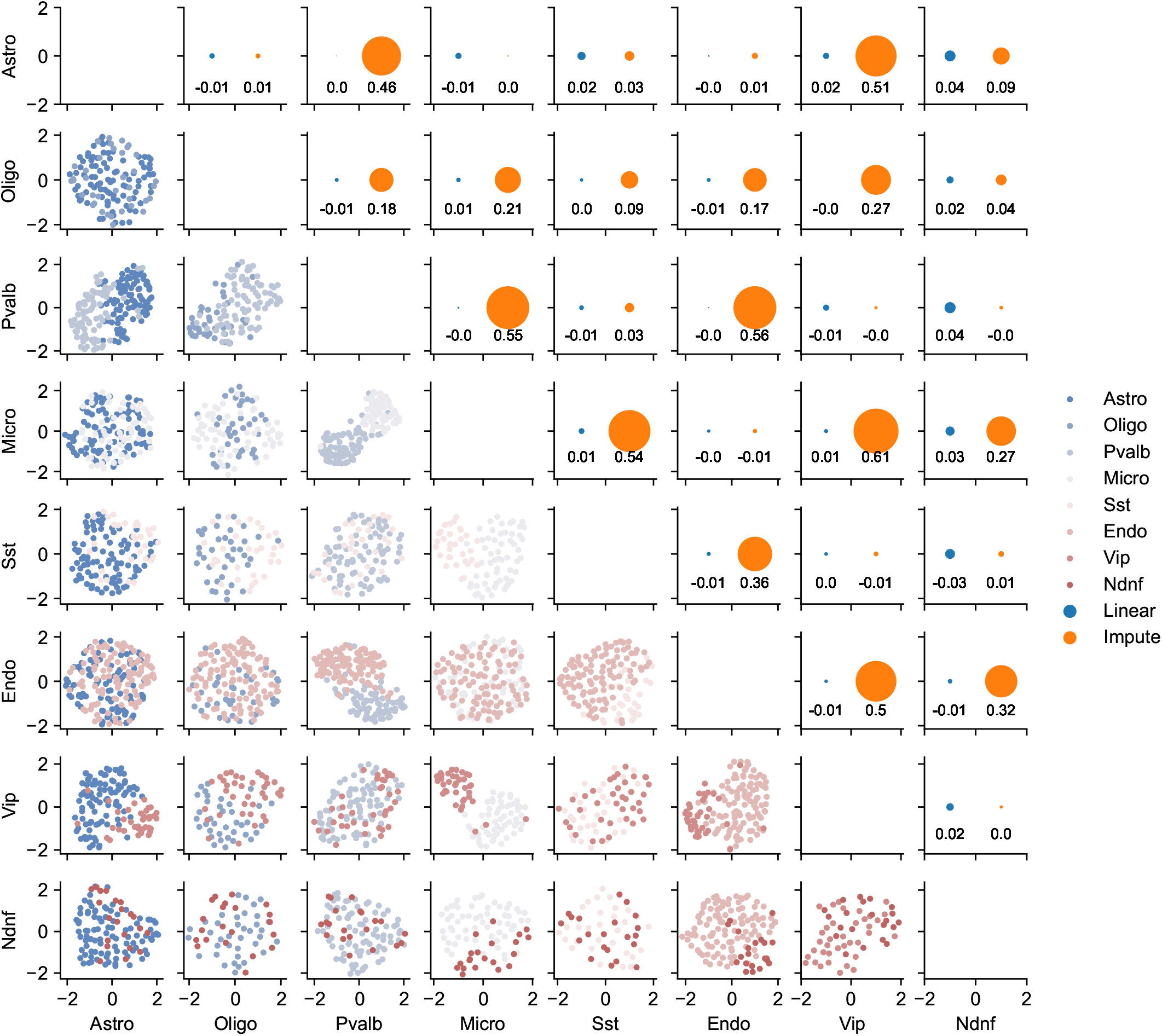
Imputation allows cell type clustering using 3D chromatin conformations. Eight major cell types of mouse brain cells are shown. Excitatory neurons are excluded because of their large numbers compared to other cell types. The lower left plots are pairwise UMAP plots between any two cell types. The dimension of the original data is first reduced to twenty with PCA before performing UMAP. The upper right plots are the adjusted mutual information scores between the clustered data and the ground truth from mRNA data. Both linear imputation result and SnapFISH-IMPUTE result are shown.

**Figure S6.**
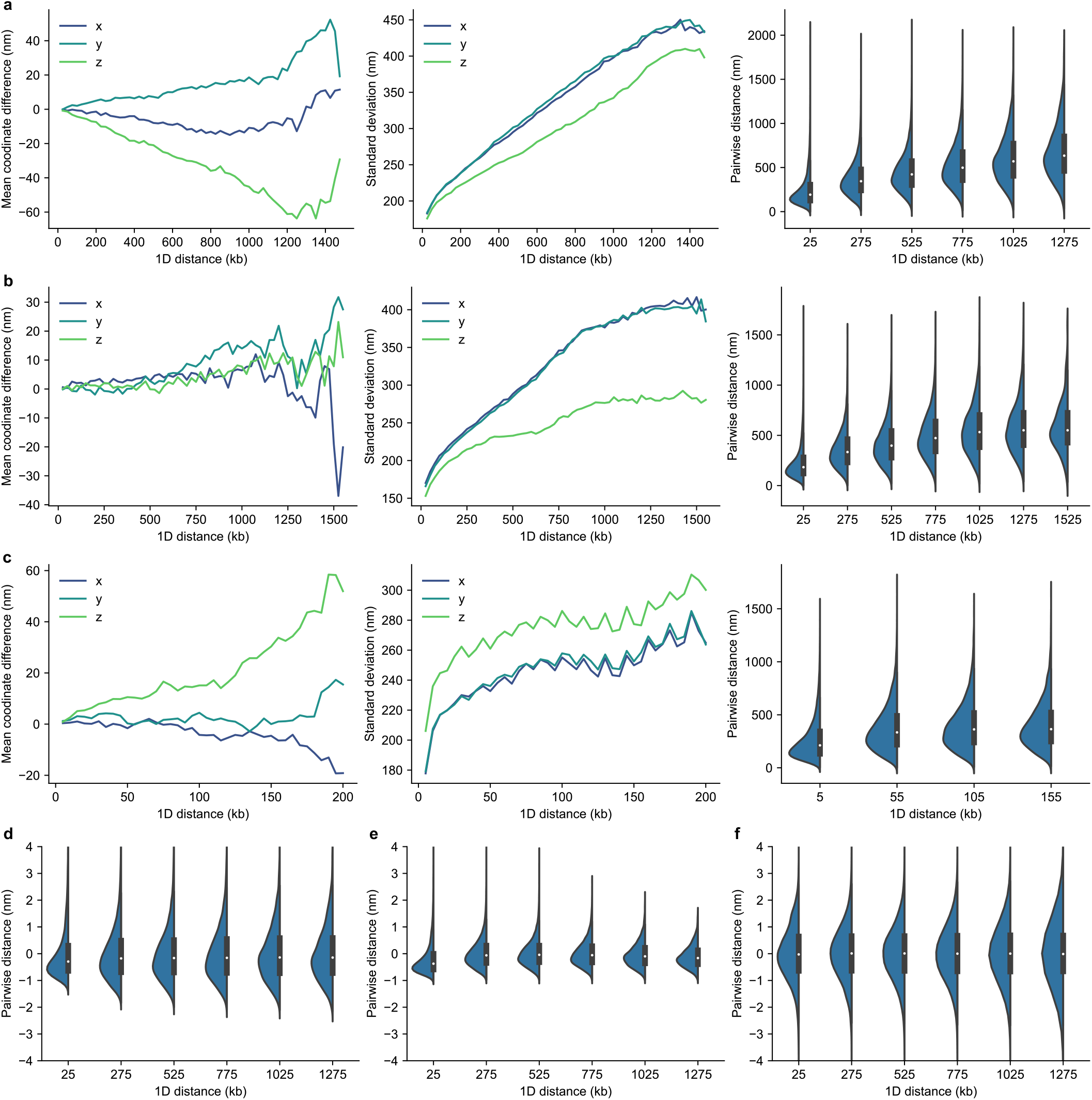
3D coordinates from different resolutions and imaging protocols have distinctive characteristics. **a** Imaging region 1 from the 25kb subset of the DNA seqFISH+ mESCs dataset. **b** Imaging region 2 from the 25kb subset of the DNA seqFISH+ mESCs dataset. **c** The 129 allele from the 5kb chromatin tracing dataset. Each row: the average distance in each axis between locus pairs with the same 1D genomic distance, the standard deviation in each axis between locus pairs with the same 1D genomic distance, and the Euclidean distances between locus pairs with the same 1D genomic distance. **d** *z*-score normalized pairwise distances. **e** GLM normalized pairwise distances. **f** Two-stage normalized pairwise distances.

## Notes

### Competing Interest Statement

The authors have declared no competing interest.

